# Engineering an anti-CD206-synNotch receptor: insights into the development of novel synthetic receptors

**DOI:** 10.1101/2024.02.07.579082

**Authors:** Sofija Semeniuk, Bin-Zhi Qian, Elise Cachat

## Abstract

Immune cells play a pivotal role in the establishment, growth and progression of tumors at primary and metastatic sites. Macrophages, in particular, play a critical role in suppressing immune responses and promoting an anti-inflammatory environment through both direct and indirect cell-cell interactions. However, our understanding of the mechanisms underlying such interactions is limited due to a lack of reliable tools for studying transient interactions between cancer cells and macrophages within the tumor microenvironment. Recent advances in mammalian synthetic biology have introduced a wide range of synthetic receptors that have been used in diverse biosensing applications. One such synthetic receptor is the synNotch receptor, which can be tailored to sense specific ligands displayed on the surface of target cells. With this study, we aimed at developing a novel *α*CD206-synNotch receptor, targeting CD206^+^ macrophages, a population of macrophages that play a crucial role in promoting metastatic seeding and persistent growth. Engineered in cancer cells and used in mouse metastasis models, such tool could help monitor and understand the effects cell-cell interactions between macrophages and cancer cells have on metastasis establishment. Here, we report the development of cancer landing pad cells for versatile applications, the engineering of *α*CD206-synNotch cells, report the measurements of their activity and specificity, and discuss the unexpected caveats when considering their *in vivo* applications.

## Introduction

The intercellular interactions, both direct and indirect, between malignant and immune cells play a significant role in cancer growth and progression[1, 2]. During all stages of cancer development through to metastasis formation, multiple subsets of immune cells can be found in the tumour microenvironment, such as cytotoxic cells (e.g., CD8^+^ T cells or NK cells), immunoregulatory Treg, Breg and T helper cells, as well as macrophages[1]. These immune cell populations contribute to cancer cell establishment and successful propagation through direct cell-cell contact or through indirect interaction via soluble cytokines[1, 3]. One of the most prominent immune cell types participating in these interactions are macrophages[2, 4, 5, 6]. In the tumour microenvironment (TME), monocytes are polarised towards either a pro-inflammatory or pro-tumorigenic state, making them an important player in tumour development and progression[4, 5, 7, 8].

Importantly, studying the processes that underlie immune cell reprogramming and cancer growth is challenging due to the transient nature of these interactions. Recent developments in synthetic biology offer receptor-based tools which allow studying various biological processes, such as tissue development[9, 10] and cell signalling[11]. The use of synthetic receptors, derived from endogenous receptors but engineered to either detect novel ligands, elicit custom responses, or both, has been largely demonstrated in the published literature and is reviewed elsewhere[11]. In the context of this study, one such tool is the synthetic Notch (synNotch) receptor, which uses synthetic input and output modules and is one of the few receptors that specifically detect membrane-tethered ligands (Fig. 1A)[10]. Multiple studies have already demonstrated the potential applications of synNotch in therapeutics and diagnostics[10, 12, 13, 14, 15, 16, 17, 18, 19, 20], tissue morphogenesis[9], and fundamental studies[21].

**Figure 1:**
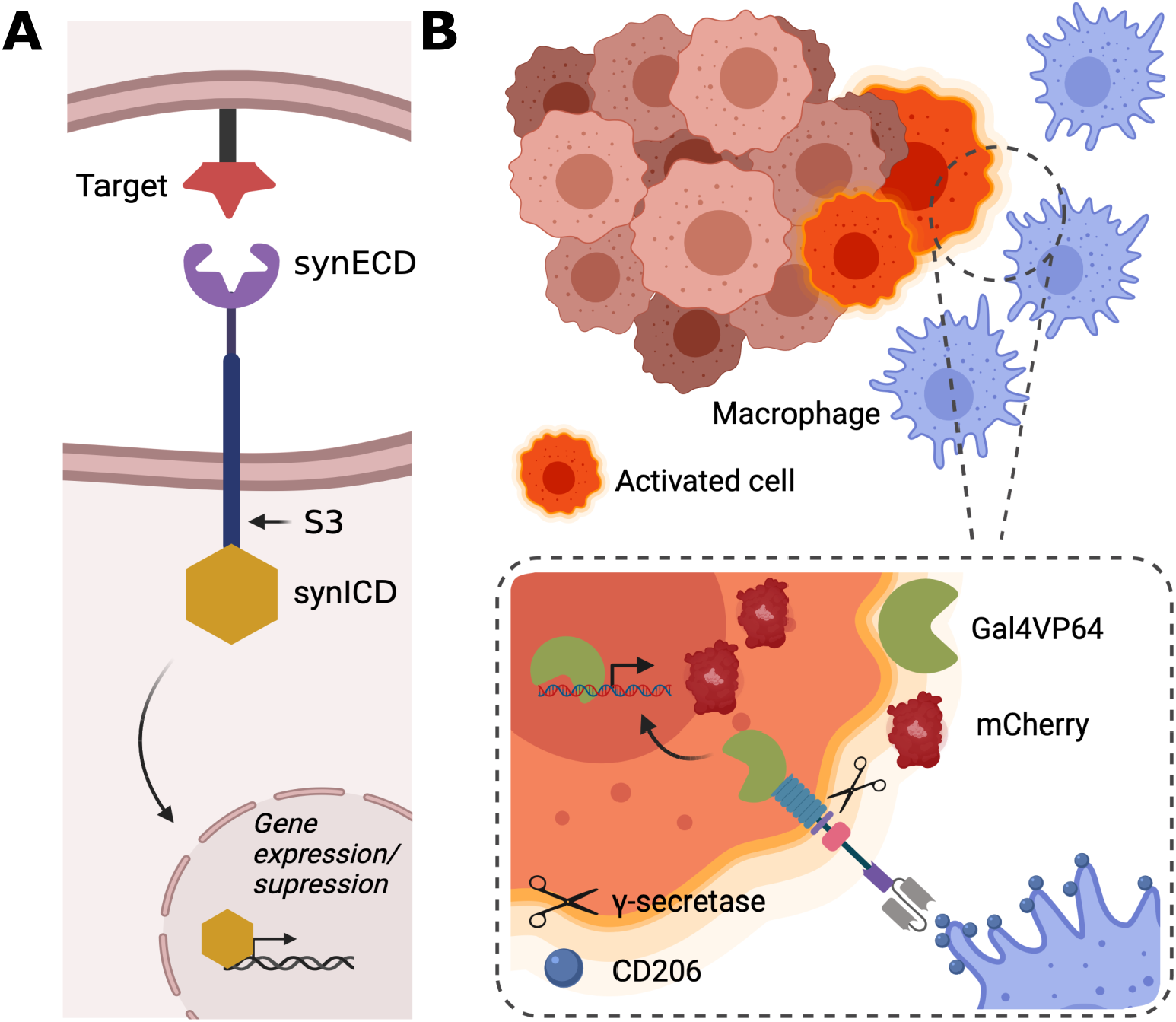
The architecture and mechanism of synNotch receptors. (A) The synNotch receptor consists of three modular domains: extracellular domain (synECD), Notch core domain and intracellular domain (synICD). S3 indicates a crucial cleavage site, which is targeted by γ-secretase. Upon ligand recognition by the synECD, mechanical forces open S3, which leads to the release of the synICD. Translocation of synICD into the nucleus can be engineered to induce changes in expression of downstream genes of interest. (B) Schematic representation of the engineered macrophage-specific synNotch system. Cancer cells engineered with the macrophage-specific synNotch detect macrophages in the tumour microenvironment. Binding between the macrophage surface marker (in this case CD206) leads to the release of transcriptional activator Gal4VP64, which translocates to the nucleus and induces the expression of a reporter gene (mCherry). *Created with BioRender.com*

We aimed to develop a synNotch-based receptor-reporter system to monitor the transient interactions between cancer cells and immune cells both *in vitro* and *in vivo* in mouse models of metastasis (Fig. 1B). Unlike other synNotch research where the focus is engineering immune cells to target cancer cells, this study aims to engineer cancer cells with a macrophage-sensitive synNotch receptor targeting CD206, a macrophage surface marker specific to pro-tumorigenic macrophage subsets. Upon ligand recognition, induction of a genetically encoded reporter results in a fluorescent response in engineered cancer cells. Extracting and sorting tumour cells into fluorescent (positive for macrophage contact) and non-fluorescent (negative for macrophage contact) cell populations will help decipher the pro-metastatic effects the cell-cell interactions between tumour and immune cells have on cancer cells and their survival, and potentially lead to the identification of new drug targets that can disrupt these effects. Here, we present the development of an anti-CD206 (*α*CD206)-synNotch receptor, together with the insights we gained from this study regarding receptor activity and specificity.

## Methods

### Molecular biology

The *α*CD206-synNotch comprised of an IgK leader peptide (derived from the Bornean orangutan T-cell surface glycoprotein CD8 alpha chain; MALPVTALLLPLALLLHAARP), myc tag, an *α*CD206 VHH sequence[22], Notch core domain and Gal4VP64 transcriptional activator[10].The *α*CD19-synNotch sequence was identical to the one published by Morsut et al.[10]. Both receptors were expressed under a mammalian phosphoglycerate kinase (PGK) promoter and had a bovine growth hormone polyadenylation (BGH polyA) sequence at the C terminal. The whole cassette was flanked by Piggy-Bac inverted terminal repeats (ITRs).

For the generation of MetBo2-CD206^+^, MetBo2-F4/80^+^ and MetBo2-CD19^+^ sender cells, CD206, F4/80 and CD19-expressing vectors were generated. The CD206 expression cassette consisted of a putative CD206 extracellular domain sequence (NM008625.2, 81 - 3835 nt), a myc tag and a PDGFRβ transmembrane domain (derived from the transmemberane domain of human platelet-derived growth factor receptor; AVGQDTQEVIVVPHSLPFKVVVISAILALVVLTIISLIILIMLWQKKPR). The CD206 sequence was extracted form IL-4 treated Bone Marrow Derived Macorphages (BMDMs). The F4/80 cassette consisted of a F4/80 coding sequence (NM010130.4, 21 – 2836 nt). The CD19 expression cassette was identical to the one published by Morsut et al.[10]. Both CD206 and CD19 cassettes were expressed under a cytomegalovirus (CMV) promoter and had a bovine growth hormone polyadenylation (BGH polyA) sequence at the C terminal.

All constructs and most essential primers used in this research are summarised in the Supplementary table 1 and 2, respectively.

### Cell culture

#### Cell lines

MetBo2 (polyoma middle T oncogene–induced mouse mammary tumor on a syngeneic Friend Virus B NIH Jackson (FVB) background)[23] cells were maintained in 1X Dulbecco’s Modified Eagle Medium (DMEM) (Thermofisher Scientific; Cat. No. 11995065) with 10 % Fetal Bovine Serum (FBS) (Sigma-Aldrich; Cat. No. F2442) and 1 % Pen/Strep (Thermofisher Scientific; Cat. No. 15140122) or Antibiotic-Antimycotic (Thermofisher Scientific; Cat. No. 15240096). cell cultures were kept at 37^*°*^ with 5 % CO_2_.

### Transfections

Cells were seeded in 48 or 24-well plates 24 h prior to transfections. For transfections, Lipofectamine 3000®(Thermofisher Scientific; Cat. No. L3000001) was used.

### Co-cultures

For co-cultures, receptor and sender cells were mixed together at a 1 : 1 ratio and seeded in a cell culture plate. For a 24-well plate format, 0.5 × 10^5^ of each cell type was used. For a 48-well format, 0.3 × 10^5^ of each cell type was used. Cells were grown in the 37^*°*^ incubator for 24 hours prior to imaging or flow cytometry.

### Flow cytometry

For the acquisition of heterogenous and monoclonal cell populations, cells were harvested 1X Accutase® (Thermofisher Scientific; Cat. No. A1110501) and centrifuged at 1000 rpm for 5 min. The pellet was resuspended in 1 ml of sorting buffer (1X DPBS, 1 % FBS, 10 % penicillin/streptomycin) and centrifuged at 500 rpm for 5 min. The pellet was resuspended again in 0.5 ml of sorting buffer and kept on ice until the sorting. FACS sorting was carried out using BD FACS Aria IIIu 4-laser/11 detector Cell Sorter (The University of Edinburgh Institute of Immunology & Infection research Flow Cytometry Core Facility). Sorted cells were seeded in a recovery medium (1X DMEM, 20 % FBS, 5 % penicillin/streptomycin).

The flow cytometry experiments were carried out using BD Fortessa with FITC, PE, PE-Dazzle, PE-Cy5, PE-CY5.5, PE Texas Red, AlexaFluor700 and BV421 filters. Cells were washed using 1X DPBS and incubated for 5 min at 37^*°*^ with 1X Accutase® (Thermofisher Scientific; Cat. No. A1110501). Cells were harvested using flow buffer (1X DPBS, 1 % FBS) and transferred to a 96-well plate for flow cytometry analysis.

In flow cytometry analysis, cells were first gated by size using forward and side scatters (SSC-A against FSC-A), and singlets were gated using forward scatters (FSC-A against FSC-W). Further gating was dependent on the type of experiment.

#### Development of the MetBo2-UAS cell line

For the assessment of MetBo2-UAS clones, the mCherry fluorescence was analysed directly following singled gating. MFI of mCherry was multiplied by the percentage of mCherry^+^ cells from the parent population (singlets) to get the total fluorescence of the cell population.

#### Development and analysis of the synNotch cell lines

Four days following transfection of MetBo2-UAS cells with the receptor cassette using the Piggy-Bac system, an initial FACS bulk sorting was carried out in order to enrich the population for BFP^+^ cells. This heterogenous population was expanded and co-cultured with CD206^+^ sender cells at a 1:1 ratio. The second round of FACS single cell sorting was carried out 24-hours post co-culture in order to isolate clones that exhibited elevated levels of mCherry fluorescence. The cells were gated by BFP fluorescence, therefore isolating only receptor cells. Subsequently, MFI of mCherry in BFP^+^ population was multiplied by the percentage of mCherry^+^ cells in the parent population of (BFP^+^) cells to get the total fluorescence of the cell population. Following the expansion of monoclonal *α*CD206-synNotch cell populations, each of them was presented to CD206^+^ sender cells at a 1:1 ratio. Receptor activation levels, indicated by elevated mCherry fluorescence, were assessed using flow cytometry through identical gating and analysis pipeline, and the clone which exhibited the highest signal-to-noise ratio was selected for further experiments.

All flow cytometry data analysis was carried out in FlowJo and GraphPad Prism.

### Immunostaining of C57BL/2 mouse spleen extract

Mouse spleen extract, pre-stained with immune-cell specific antibodies was acquired from the Binzhi Qian lab at the MRC Centre for Reproductive Health at The University of Edinburgh. The following antibodies were used: Alexa Fluor® 700 anti-mouse F4/80 Antibody (Biolegend; Cat. No. 123129), PE anti-mouse CD206 (MMR) Antibody (Biolegend; Cat. No. 141705), PE/Cyanine5 anti-mouse/human CD45R/B220 Antibody (Biolegend; Cat. No. 103209), PE/DazzleTM 594 anti-mouse CD3 Antibody (Biolegend; Cat. No. 100245), PerCP/Cyanine5.5 anti-mouse/human CD11b Antibody (Biolegend; Cat. No. 101227). The extract was split into equal parts and 250 to 500 μl of supernatant, containing the small antibody domain chromobodies were loaded on the extract and incubated in the dark for 1h at 4^*°*^C. Next, the cells were twice washed with DPBS and analysed using flow cytometry. Compensation was carried out using UltraComp eBeads™ Compensation Beads (Invitrogen; Cat. No. 01-2222-42).

### Chromobody staining

Chromobodies were generated by transiently expressing the chromobody expressing plasmids in HEK293FT cells in a 6 well plate. The media was collected from the cells two days later, centrifuged to pellet the cells and cell debris. The supernatant was used to stain the cells at 4*°*C overnight in the dark.

### Fluorescent microscopy

Fluorescent imaging was carried out using Leica DMi8 fluorescent microscope with DAPI (Ex: 350/50, Em: 460/50), TexasRed (Ex: 560/40, Em: 630/75), GFP (Ex: 470/40, Em: 525/50) and Y5 (Ex: 620/60, Em: 700/75) filter cubes. Further image processing was carried out in FIJI software.

## Results

### Development of a stable *α*CD206 synNotch cell line

We sought to achieve stable genomic integration of the synNotch system in MetBo2 cells, a bone metastasis cell line derived from mouse mammary tumour background[23]. We chose ROSA26 safe harbour[24] for the integration of the reporter cassette. The strategy was adapted from Malaguti et al. (Fig. 2A)[25]. First, a landing pad was established using CRISPR/Cas9-guided integration through homology-directed repair (HDR). The landing pad consisted of a nuclear mKate2 expression cassette (CAG-mKate2-3xNLS), with an upstream promotorless Neomycin resistance (NeoR) open reading frame (ORF), expressed exclusively upon correct targetting of the construct downstream of the ROSA26 endogenous promoter. The whole landing pad cassette was flanked with the attP50 recombination sites for later Φc31-mediated cassette exchange. Following antibiotic selection and clonal isolation of mKate2^+^ cells, the UAS-mCherry reporter cassette was integrated through Φc31-mediated cassette exchange (RMCE). The reporter cassette consisted of a puromycin resistance (Pac) ORF at the 5’ of the UAS-mCherry cassette. Following RMCE and antibiotic selection, the now mKate2^-^ cells were sorted into single cells. Expanded monoclonal cell populations were tested for activation upon transfection with Gal4VP64 transcriptional activator, and the best performing MetBo2-UAS clone (317.1-fold activation) was chosen for further experiments (Fig. 2B, C). Genomic integration into the mROSA26 safe harbour was also validated through PCR on genomic DNA (Fig. 2D) The *α*-CD206 synNotch receptor cassette was integrated in MetBo2-UAS cells through PiggyBac transposase-based integration. The receptor architecture consisted of a *α*CD206 nanobody [22], fused to the Notch core domain and a Gal4VP64 transcriptional activator (Fig. 2E). Downstream from the receptor cassette we integrated a lineage tracking component - H2B-BFP cassette - which was used as selection marker throughout the first round of FACS sorting. The whole receptor and H2B-BFP construct was flanked by PiggyBac inverted terminal repeats (ITRs). The detailed description of cell line development methodology is available in the Methods section.

**Figure 2:**
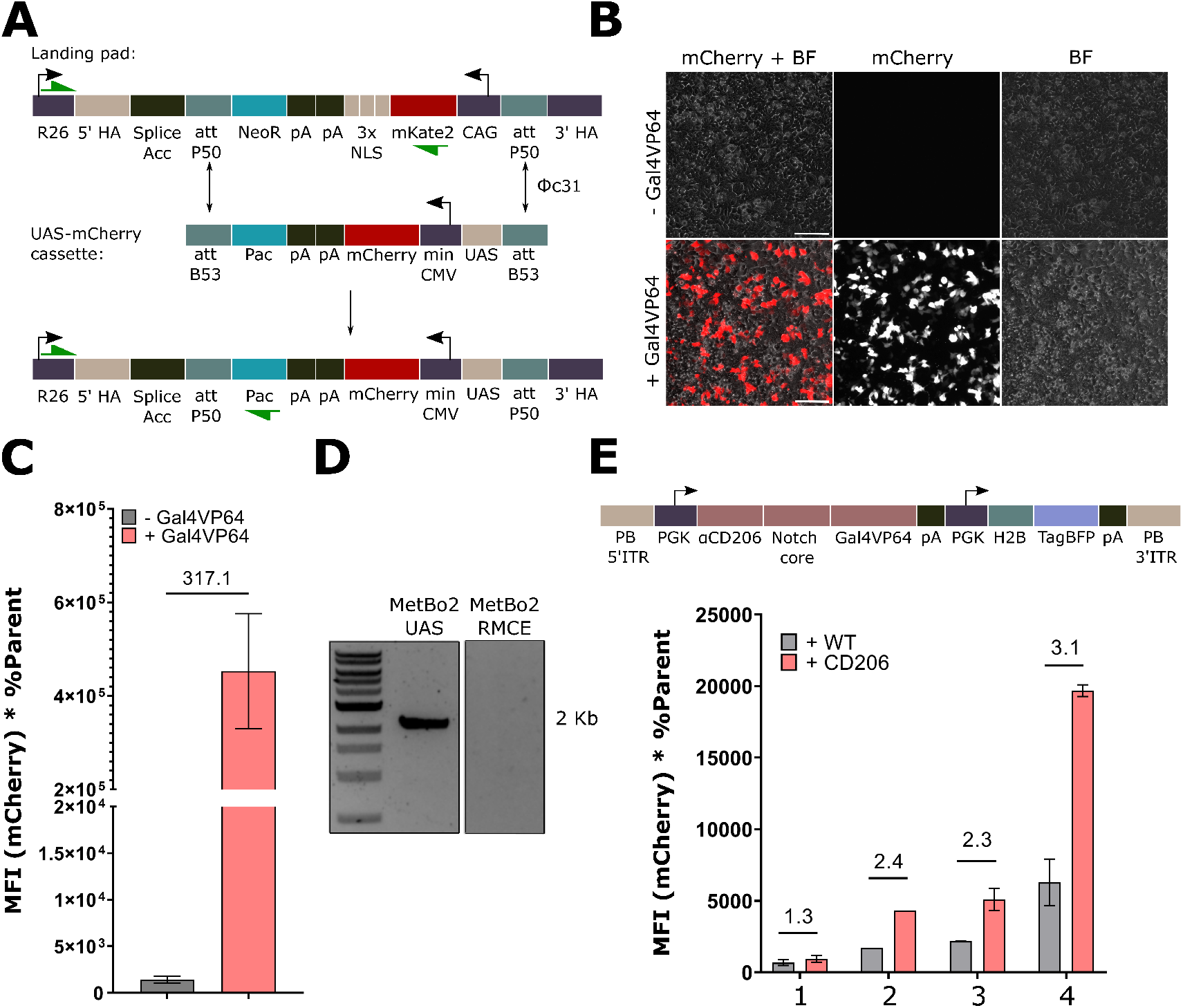
Engineering of the *α*CD206-synNotch cells. (A) Design of the MetBo2-UAS cell line. Initially, a landing pad was established, comprising two selection markers: a G418 resistance cassette, activated by the endogenous ROSA26 promoter upon successful integration, and a nuclear mKate2 cassette. Subsequently, through Φc31 recombinase, this cassette was exchanged for a minCMV-UAS-mCherry cassette also replacing the neomycin resistance cassette for a puromycin one. (B, C) Selected MetBo2-UAS clone exhibited inducible (317.1-fold) mCherry fluorescence upon transfection with Gal4VP64. Scale bar 50 μm. (D) Integration into mROSA26 locus was confirmed by PCR of genomic DNA (2,175 bp). (E) Development of the *α*CD206-synNotch construct. Four clones of the *α*CD206-synNotch receptor were isolated and tested for activation with CD206^+^ cells. All clones were analysed in triplicates except for the clone 2. *Green arrows indicate primer binding sites. HA – homology arms. Kan/NeoR – Kanamycin/Neomycin (G418) resistance gene. pA – polyadenylation sequence. NLS – nuclear localisation sequence. CAG – Cytomegalvirus immediate enhancer/*β*-actin promoter. Pac – puromycin acetylase (puromycin resistance gene). PGK – phosphoglycerate kinase promoter. scFV – single chain variable fragment. H2B – human histone 2B*.

Following co-culture screening of the *α*-CD206 synNotch clone candidates, four monoclonal populations of *α*CD206-synNotch Metbo2 cells were isolated (Fig. 2E). Clone number 4 (3.1-fold activation) was chosen as the best-performing clone when tested against CD206^+^ sender cells (see below) and will be further referred to as *α*CD206-synNotch.

### Development of CD206^+^ sender cells

We engineered synthetic CD206^+^ sender cells expressing the extracellular domain (ECD) of mouse CD206. The CD206 expression cassette contained an ORF for the CD206 ECD (NM008625.2, 81 - 3835 nt) fused to the PDGFRβ transmembrane domain (Fig. 3A). The CD206 ECD sequence was isolated from the cDNA of BV6 mouse bone marrow-derived macrophages (BMDMs) following their induction with interleukin-4 (Fig. 3A). In this proof-of-concept study, synthetic sender cells were preferred over primary macrophages due to sourcing issues. The whole cassette was transiently expressed in sender cells and validated through immunostaining using *α*CD206-mNeonGreen chromobodies in an assay developed by Baronaite et al. (Fig. 3B)[26] (See Methods). Specifically MetBo2 cells were chosen as the sender cell chassis to minimise the possibility of ligand-independent receptor activation and/or receptor cis-activation from non-canonical ligands present on the surface of MetBo2 cells within the downstream co-cultures due to cell-cell interactions between different cell lines or types.

**Figure 3:**
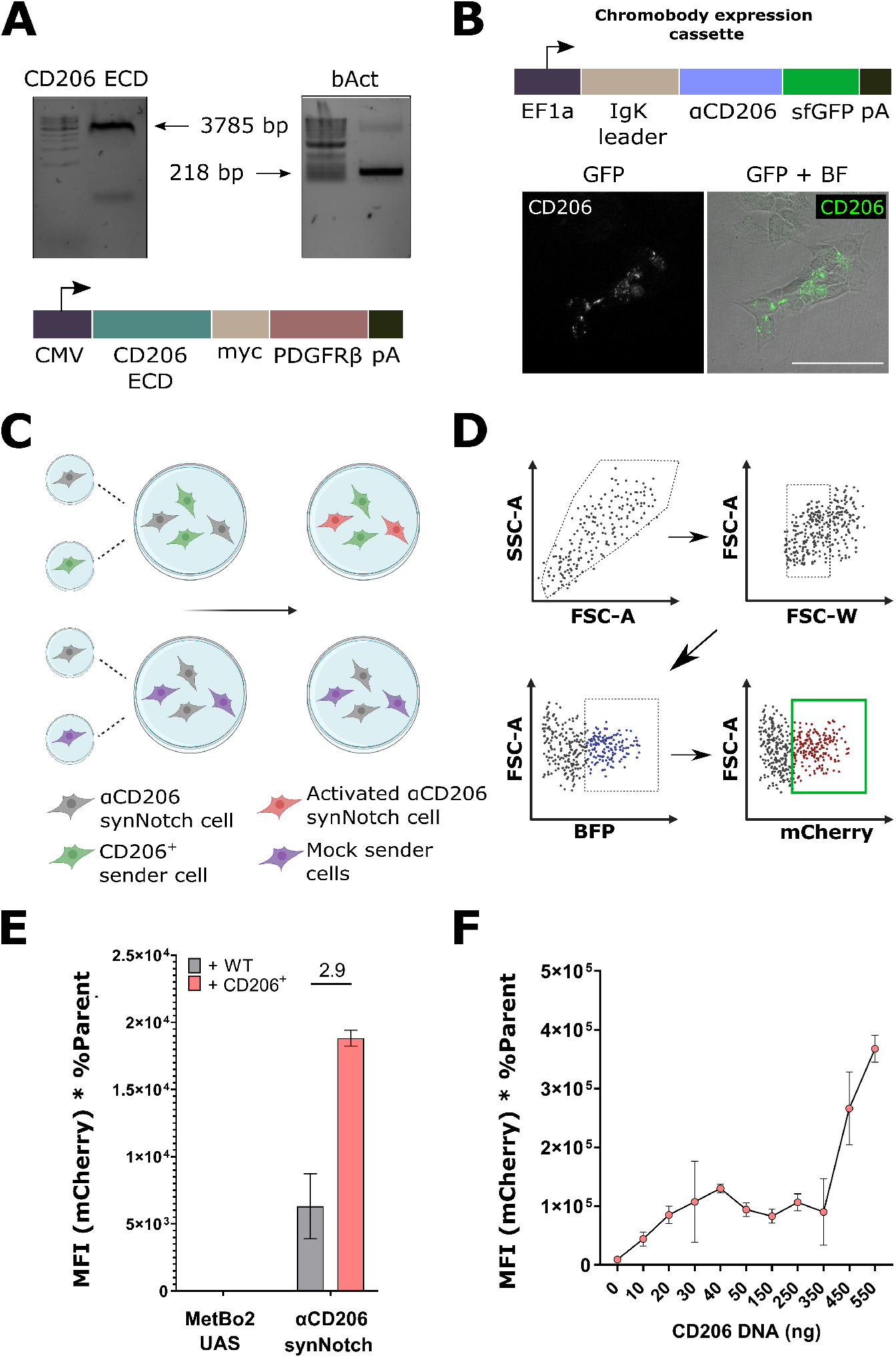
Development and validation of an *α*CD206-synNotch receptor. (A) Extraction of CD206 ECD CDS from cDNA of IL4 stimulated bone marrow-derived mouse macrophages (BMDM). *β*-actin was used as housekeeping gene for validation of cDNA. (B) Positive staining by *α*CD206 chromobodies (GFP) of live MetBo2 sender cells transiently expressing the CD206 ECD construct. Scale bar 100 μm. (C) Co-culture strategy for normalisation of co-culture conditions among test and control wells. *Created with BioRender.com* (D) Flow cytometry gating strategy to quantify the mCherry fluorescence of synNotch-positive (BFP^+^) activated cells. *Created with BioRender.com* (E) *α*CD206-synNotch exhibits a 2.9-fold activation in co-culture with CD206^+^ cells. In comparison, no mCherry signal was observed when using MetBo2-UAS cells as receiver cells, which shows that in synNotch co-cultures the mCherry signal comes solely from receptor activation. (F) *α*CD206-synNotch demonstrates an increase in activity in response to increasing amounts of ligand transfected in sender cells. *CMV – Cytomegalovirus mammalian promoter. HA - Human influenza hemagglutinin tag. Myc – c-myc tag. PDGFR*β *- Platelet-derived growth factor receptor beta trans-membrane domain. pA – polyadenylation sequence*.

### *α*CD206-synNotch successfully targeted CD206^+^ cells *in vitro*

The co-culture strategy used to determine receptor activation is depicted in figure 3C. Here, *α*CD206 synNotch cells were co-cultured either with ligand-presenting sender cells (CD206^+^ cells), or mock sender cells (wild-type MetBo2 cells). This allowed to normalise receptor activation between test and control groups by maintaining equal numbers of receiver to sender cells in co-cultures. Additionally, our flow cytometry gating strategy to quantify the percentage of activated cells is depicted in figure 3D. Here, BFP is associated with the constitutive H2B-BFP expression from the receptor cell population, and mCherry is the reporter expressed upon contact with sender cells and marker of activated cells. The best performing *α*CD206-synNotch clone from the initial screen was re-tested for activation and exhibited a 2.9-fold activation when co-cultured with MetBo2-CD206^+^ cells (Fig. 3E). Moreover, the clone also demonstrated a dose-dependent activation pattern, with a sharp increase in activation when the sender cells were transfected with more than 450 ng of the ligand expressing vector in a 48 well plate (Fig. 3F).

### *α*CD206 synNotch exhibits cross-specificity with other ligands

To better characterise and assess the suitability of *α*CD206-synNotch in preparation for *in vivo* applications and the targeting of CD206^+^ macrophages, we tested the *α*CD206-synNotch cells for cross-reactivity with cells overexpressing an irrelevant surface ligand: human CD19, a distinct marker of B cells and a commonly used ligand in other synNotch applications [10, 13, 12, 19]. We engineered a human CD19[10] expression vector (Fig. S1A) and generated MetBo2-CD19^+^ sender cells (Fig. S1B) for co-culture experiments in parallel with CD206 sender cells. Interestingly, *α*CD206-synNotch exhibited activation (22.2-fold increase) when co-cultured with CD19^+^ sender cells (Fig. 4A). Moreover, we observed significant fluctuations in synNotch activity over passages, with *α*CD206-synNotch activation levels reaching 23.6-fold, compared to the 2.9-fold activation measured previously. The possible reasons for fluctuations in the receptor activation are considered in the Discussion section. For comparison, we tested the *α*CD19-synNotch[10] architecture for reciprocal cross-reactivity with MetBo2-CD206^+^ sender cells and didn’t observe any.

**Figure 4:**
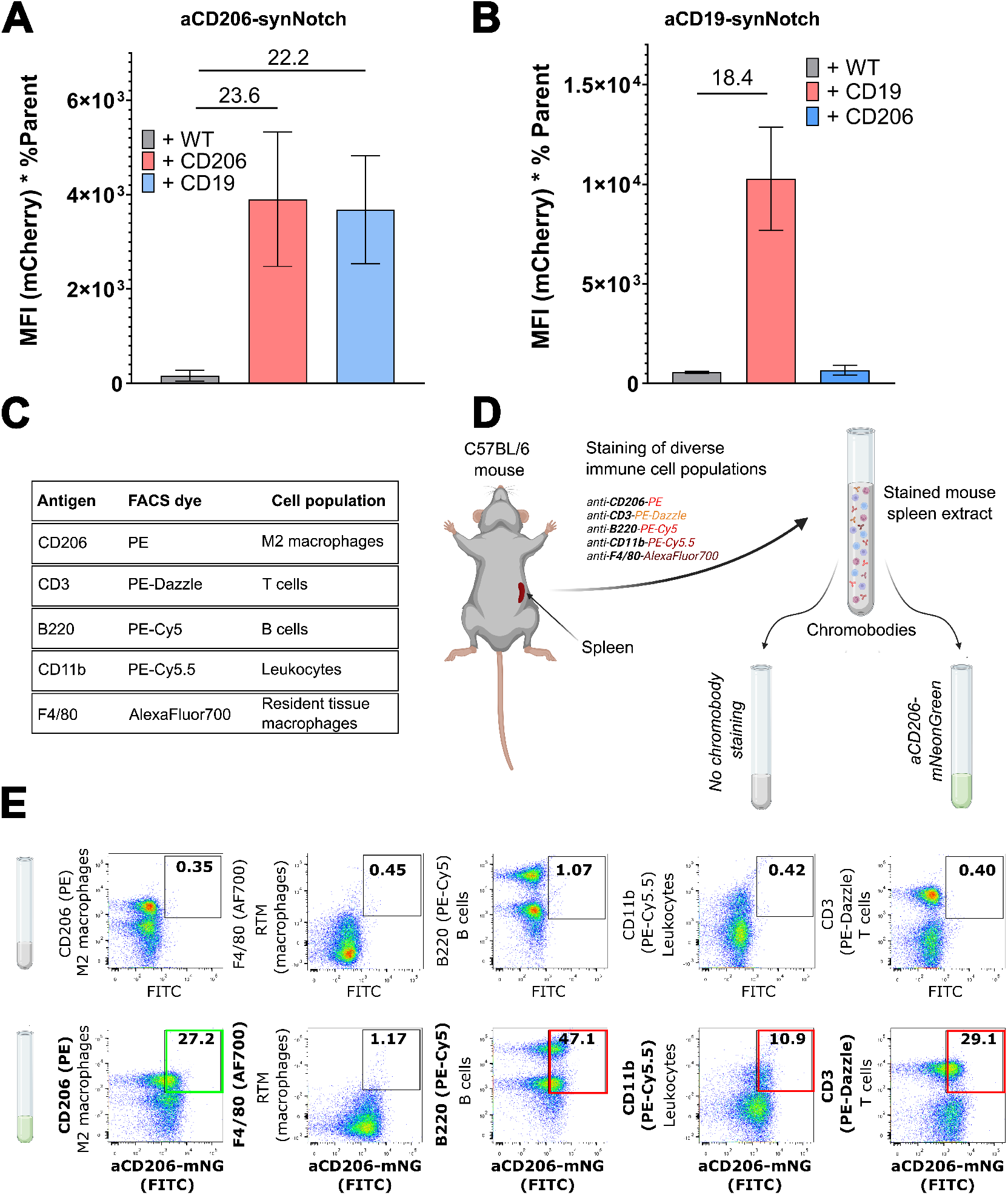
Testing the specificity of the *α*CD206 synNotch. (A) *α*CD206-synNotch cells exhibited significant cross-reactivity when presented to CD19^+^ cells. (B)*α*CD19-synNotch cells exhibited no cross-reactivity when co-cultured with CD206^+^ sender cells. (C) The list of antibody antigens and corresponding conjugated dyes used to stain the C57BL/6 mouse spleen extract for specific immune cell subpopulations. (D) The mouse spleen extract was stained with a mix of antibody conjugates in a single-pot reaction. Equal parts of the mix were then stained with *α*CD206-VHH fused to mNeon-Green (chromobodies). *Created with BioRender.com* (E) Flow cytometry evaluation of co-staining with the various immune cell subpopulation specific marker dyes: (top) no chromobody co-staining, and (bottom) co-staining with *α*CD206-mNeonGreen. Numbers indicate the percentage of the co-stained populations.

To assess whether synNotch cross-reactivity is due to the low specificity of the small antibody domains used as synNotch extracellular domains (ECDs), we tested the *α*CD206 VHH for cross-reactivity against various endogenously expressed murine immune cell markers. Through our testing platform, we evaluated the affinity of *α*CD206-mNeonGreen chromobodies for various immune cell types from a C57BL/6 mouse spleen extract. First, this spleen extract was stained with five conjugated antibodies specific to distinct immune murine cell markers: CD206 (Phycoerythrin, PE), CD3 (PE-Dazzle), B220 (or CD45, PE-Cy5), CD11b (PE-Cy5.5) and F4/80 (AlexaFluor700) (Fig. 4C, D). These markers corresponded to five different immune cell populations: CD206^+^ pro-inflammatory macrophages, T cells, B cells, leukocytes and F4/80^+^ resident tissue macrophages, respectively. The dye-stained extract was then cross-stained with *α*CD206-mNeonGreen chromobodies. The *α*CD206 VHH reacted with CD206^+^ M2 macrophages as expected, but cross-reacted with all other tested immune cell populations: CD3^+^ T cells, B220^+^ B cells and CD11b^+^ macrophages, showing poor specificity for its cognate target.

## Discussion

In recent years the use of synNotch receptors has been widely demonstrated in a variety of applications, both *in vitro* and *in vivo*[10, 13, 12, 14, 19, 20, 25, 18, 9, 21, 16, 18]. The majority of such applications are within the field of cancer oncology where the synNotch has been employed to target and eliminate malignant cells. However, none of these research applications have, to our knowledge, reported any receptor cross-specificity. Here, we demonstrated that a newly developed synNotch receptor can exhibit cross-specificity with other cell surface markers.

The cross-reactivity of small antibody domains used as synNotch ECDs with various immune cell populations was evaluated using a mouse spleen extract as a pool of immune cells presenting various surface markers. Our findings suggest that applying the synNotch system *in vivo* presents significant challenges due to the potential activation of synNotch cells by incorrect interaction partners, resulting in false positives. For instance, while using *α*CD206-synNotch to target CD206^+^ macrophages, the receptor is likely to report cell contact with B cells (CD19^+^), which are abundant both at the primary tumour and metastatic sites[1]. Moreover, the activation of synNotch reporter cells by multiple non-target immune cells is likely to occur shortly after injection while circulating in the bloodstream, prior to the establishment of primary and, subsequently, metastatic tumors in mice. This is a crucial caveat for *in vivo* applications of these receptor-based systems, due to the possibility of false-positive detection events, as well as off-target events that may result in significant side effects in cell therapy contexts [27]. While the *α*CD206-synNotch was developed primarily to study cell interactions in a mouse model, there are many synNotch- and other synthetic receptor-based tools being developed for human cell therapy applications[12, 14, 17, 16, 18]. Therefore, we strongly suggest cross-specificity tests to be routinely implemented as a vital part of synthetic receptor development pipelines. Collectively, these findings demonstrate that the development of high-specificity nanobodies and single chain variable fragments is crucial to improving the reliability and safety of synthetic receptor systems.

Other measurements to minimise the detection of false-positives and mitigating the effect of receptor cross-reactivity are to (i) evaluate the duration of synNotch activation and (ii) evaluate the ligand exposure time needed to induce the fluorescent response. Knowing the amount of time needed for the cell-cell interaction to induce a fluorescent response, as well as knowing the response duration, would allow for more precise temporal discrimination between false positive activation and ligand-specific activation. Alternatively, resorting to partially immunodeficient mouse strains may help to reduce the amount of non-specific receptor activation[28]. However, this approach is limited to certain applications, as these tools are specifically developed to track interactions with immune cells.

Lastly, we obeserved that *α*CD206-synNotch exhibited a variable and non-reproducible pattern of activation levels, i.e. an inconsistent increase in mCherry fluorescence levels in co-cultures with MetBo2-CD206^+^ cells over cell passages. This variability might stem from the use of transient transfection for the generation of sender cells, which resulted in variable levels of ligands, despite the fact that the same cell numbers and growing conditions were applied throughout all co-culture experiments. Therefore, stable ligand-expressing sender cells should be used in order to ensure more consistent co-culture conditions.

Collectively, these findings indicate that utilising synNotch, as well as other synthetic receptor-based systems *in vivo* presents potential risks related to receptor cross-reactivity. Therefore, such tools and their applications must be properly characterised and validated by incorporating cross-specificity tests into the standardised receptor testing pipelines.

## Acknowledgements

We would like to thank Weijia Liu, Dr. Vivek Senthivel, Ugne Baronaite, Huanwen Wu, Dr. Ruoyu Ma, Chengbin Zhang, Dr. Anu Fernando and other members of Elise Cachat, Bin-Zhi Qian and Susan Rosser laboratories. Sanger sequencing was carried out by Edinburgh Genomics at the University of Edinburgh. Edinburgh Genomics is partly supported through core grants from NERC (R8/H10/56), MRC (MR/K001744/1) and BBSRC (BB/J004243/1). Flow cytometry data were generated within the Flow Cytometry and Cell Sorting Facility in Ashworth, King’s Buildings at the University of Edinburgh. The facility is supported by funding from Wellcome and the University of Edinburgh. Funding: This work was supported by the EPSRC (EP/R513209/1), the Wellcome-University of Edinburgh ISSF3 fund (IS3-R69), the School of Biological Sciences at the University of Edinburgh. Funding: This work was supported by the EPSRC (EP/R513209/1), the Wellcome-University of Edinburgh ISSF3 fund (IS3-R69), the School of Biological Sciences at the University of Edinburgh.

