## Supplementary Information for "Engineering an anti-CD206-synNotch receptor: insights into the development of novel synthetic receptors"

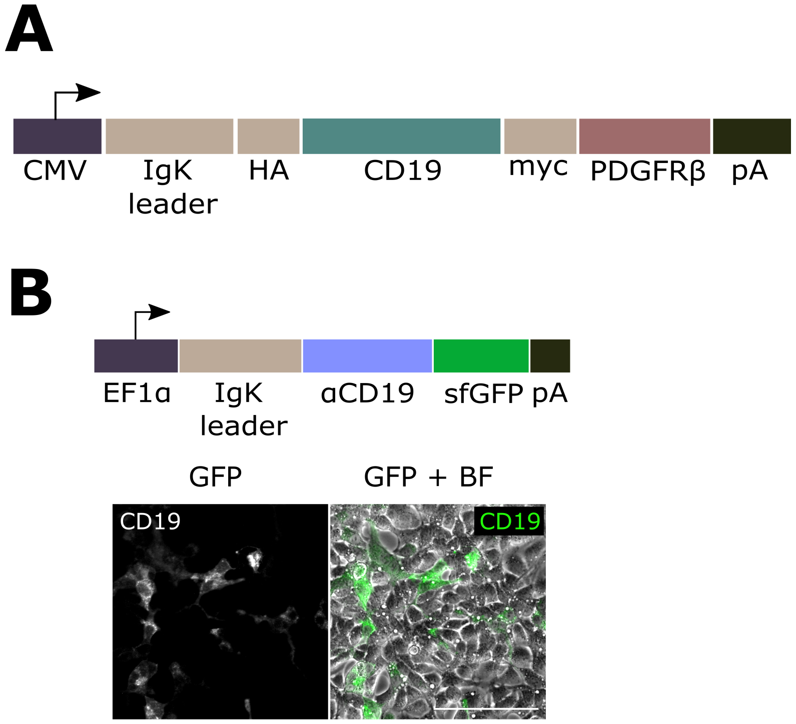


**Supplementary Figure 1: Development of MetBo2-CD19⁺ sender cells.** (A) Structure of the CD19 construct. (B) Chromobody staining of MetBo2 cells, transiently transfected with CD19 surface ligand. Positive staining is indicated by GFP fluorescence. Scale bar 100 μm.

**Supplementary Table 1: Summary of the constructs.**

| Name | Description | Construct | Notes |
|---|---|---|---|
| Addgene #79125 | $\alpha$CD19-synNotch | PGK→IgKleader→myc→$\alpha$CD19→ synNotch→ Gal4VP64→WPRE | A gift from Wendell Lim[10] |
| KPL155 | $\Phi$c31 recombinase | CMV→$\Phi$c31→pA | A gift from Sally Lowell[25] |
| Addgene #183609 | ROSA26 landing pad for creation of MetBo2-RMCE | ROSA HA 5'→Splice Acc→ Kan/NeoR→pA←pA←3xNLS-mKate2←CAG←ROSA HA 3' | A gift from Sally Lowell[25] |
| pHWu1 | $\alpha$CD206- synNotch | PGK→IgKleader→6xHis→ $\alpha$CD206→synNotch→ Gal4VP64→WPRE | |
| pSSe3 | CD19 ligand for engineering of MetBo2 CD19$^{+}$ cells | CMV→IgKleader→HA→CD19→ myc→PDGFRβ→pA | |
| pSSe14 | CD206 ligand for engineering of MetBo2 CD206$^{+}$ cells | CMV→CD206→myc→PDGFRβ→ pA | |
| pSSe22 | $\alpha$CD19-synNotch | PB ITR 5'→PGK→IgKleader →myc→$\alpha$CD19→synNotch→ Gal4VP64→pA→PGK→H2B-TagBFP → pA →3' PB ITR | PiggyBac backbone |
| pSSe24 | $\alpha$CD206-synNotch | PB ITR 5'→PGK→IgKleader →myc→$\alpha$CD206→synNotch→ Gal4VP64→pA→PGK→H2B-TagBFP→pA →3' PB ITR | PiggyBac backbone |
| SP59 | PiggyBac transposase | CMV→hyPBase→pA | |
| pSSe40 | UAS-mCherry cassette for RMCE | attB53→Pac→pA→pA←mCherry← minCMV←5xGal4-UAS←attB53 | |
| pSSe59 | ROSA26 gRNA and Cas9 vector | U6→gRNA→CMV→3xFLAG-Cas9-T2A-GFP→pA | |
| UBa1006 | $\alpha$CD206 VHH fused to mNeonGreen | EF1$\alpha$→IgKleader→$\alpha$CD206-mNeonGreen→pA | Engineered by Ugne Baronaite (Cachat lab) |
| UBa1007 | $\alpha$CD19 scFV fused to mNeonGreen | EF1$\alpha$→IgKleader→$\alpha$CD19-mNeonGreen→TCS-6xHis→pA | Engineered by Ugne Baronaite (Cachat lab) |

**Supplementary Table 2: Summary of the key primers**

| Name | Sequence | Notes |
|---|---|---|
| mROSAwtF | GGCGGACTGGCGGGACTA | Wild-type ROSA26 locus fwd primer; Used for confirming integration with gRNA PCR; |
| PuroR | CTTCCATCTGTTGCTGCG | Specific to puromycin resistance gene; Used as a reverse primer for confirming the integration of UAS-mCherry cassette with gDNA PCR; |
| mKateR | TACGAAGACGGGGGCGTGC | mKate2 reverse primer; Used for confirming integration with gDNA PCR; |
| mActB_F<br>mActB_R | CTGTCCCTGTATGCCTCTG<br>ATGTCACGCACGATTTTC | Murine β-actin primers used as control during extraction of ligands from cDNA; |
| MRC1_F<br>MRC_Rfull | CCGCCAGTGTGCTGGAATTCGGAAGATCCACTCTGGGCC<br>ATGAGTTTTTGTTCGTCGACGCCATAGAAAGGAATCCACGC | CD206 Gibson primers for extraction from cDNA. |
| Gibson_CD19_F<br>Gibson_CD19_R | CCCAGCCGGCCAGATCTCCCGAGGAACCTCTAGTG<br>GATGAGTTTTTGTTCGTCGACCTTCCAGCCACCAG | CD19 Gibson primers for extraction from cDNA. |
